# Information redundancy across spatial scales modulates early visual cortical processing

**DOI:** 10.1101/2021.06.29.449223

**Authors:** Kirsten Petras, Sanne ten Oever, Sarang S. Dalal, Valerie Goffaux

**Affiliations:** Psychological Sciences Research Institute (IPSY), UC Louvain, Belgium; Institute of Neuroscience (IONS), UC Louvain, Belgium; Department of Cognitive Neuroscience, Maastricht University, The Netherlands; Max Planck Institute for Psycholinguistics, The Netherlands; Donders Institute for Cognitive Neuroimaging, Radboud University, The Netherlands; Center of Functionally Integrative Neuroscience, Aarhus University, Denmark

## Abstract

Visual images contain redundant information across spatial scales where low spatial frequency contrast is informative towards the location and likely content of high spatial frequency detail. Previous research suggests that the visual system makes use of those redundancies to facilitate efficient processing. In this framework, a fast, initial analysis of low-spatial frequency (LSF) information guides the slower and later processing of high spatial frequency (HSF) detail. Here, we used multivariate classification as well as time-frequency analysis of MEG responses to the viewing of intact and phase scrambled images of human faces to demonstrate that the availability of redundant LSF information, as found in broadband intact images, correlates with a reduction in HSF representational dominance in both early and higher-level visual areas as well as a reduction of gamma-band power in early visual cortex. Our results indicate that the cross spatial frequency information redundancy that can be found in all natural images might be a driving factor in the efficient integration of fine image details.

## Introduction

In natural images, low and high spatial frequencies are spatially aligned such that the low spatial frequency (LSF) phase i.e., the location of low spatial frequency features, is informative towards the content and location of high spatial frequency (HSF) contrast. This alignment of spatial frequencies is essential for efficient visual perception (Bex and Makous, 2002; MaBouDi *et al.*, 2016).

Efficient coding theories (Barlow, 1961, 2001; Simoncelli, 2003) conjecture that the visual system aims to exploit redundancies in the form of statistical dependencies across image scales to optimize processing. The initial assessment of coarse image structure and semantic content in higher-level areas has been proposed to subserve a (predictive) feedback signal, which restricts low-level processing to the HSF details that are most relevant for recognition (Bullier, 2001; Bar *et al.*, 2006), therefore alleviating the low-level computational resources recruited for the processing of HSF. This framework is supported by empirical evidence showing that LSF is processed before HSF image content. Human object recognition is often found to be characterized by a low spatial frequency (LSF) bias at the earliest latencies of visual processing (Parker and Dutch, 1987; e.g., Mazer *et al.*, 2002) and direct comparisons of visual response times to LSF and HSF stimuli indicate a temporal advantage for LSF processing (Vlamings, Goffaux and Kemner, 2009; Goffaux *et al.*, 2010; Musel *et al.*, 2014). However, because past research usually split natural images into separate SF bands to study their respective contribution to perception, it failed to test a core assumption of the coarse-to-fine framework, namely that, in natural broadband stimulation, the early encoding of the coarse LSF reduces the computational resources recruited for the subsequent processing of HSF details. This assumption can indeed only be tested in broadband viewing conditions, containing phase-aligned (redundant) coarse and fine input.

In a recent work, we developed a novel approach to track the separate contribution of LSF and HSF processing to the human perception of *broadband* images of faces. In this approach, multivariate pattern classifiers are first trained to differentiate between human EEG scalp amplitude patterns elicited by LSF and HSF stimuli devoid of any semantic content (i.e., phase-scrambled images), and tested on how they generalize to broadband visual stimulation over time. Broadband images contain both LSF and HSF information, so classifier generalization indicates which profile is most prominent in the activity pattern they elicit. Using this approach, we found that LSF/HSF classifiers tested on the neural responses to broadband images were less likely to classify responses as HSF when images showed intact faces, compared to scrambled faces. Our findings are in line with the hypothesis that LSF-driven guidance of HSF processing results in an overall reduction of processing load, when coarse (LSF) image contrast is aligned and thus informative towards its fine (HSF) content (Petras *et al.*, 2019). However, our previous methodology was limited to reveal *relative* changes in HSF, compared to LSF markers in intact vs. scrambled conditions. Whether such relative reductions in HSF dominance reflect an *absolute* reduction in the resources recruited for the processing of broadband, SF-redundant, stimulation is unclear. Furthermore, the poor spatial resolution of the employed EEG technique precludes the localization of the observed modulations.

Here, we exploit the superior spatial and temporal resolution afforded by magnetoencephalography (MEG) and explicitly test the hypothesis that coarse LSF image structure guides the processing of redundant HSF detail in early visual cortex, therewith reducing local processing load. In a further step, we investigate the proposal that such LSF facilitation is driven by feedback from higher level visual areas. Using source reconstruction, we separate the responses of high- and low-level visual regions and investigate their dynamic interactions during the processing of broadband images of faces. Electrophysiological studies in the primate visual system found that feedforward/feedback communication employs distinct response temporal dynamics. Feedforward signals leaving the superficial (supragranular) layers of cerebral cortex to target layer 4 of upstream regions are carried by fast, gamma-range (>30 Hz), neural oscillations. In contrast, feedback signals, mostly stemming from deep (infragranular) layers manifest as slow neural oscillations in the alpha and/or beta ranges (Bastos *et al.*, 2015; Fries, 2015; Michalareas *et al.*, 2016) that are primarily inhibitory in nature (Thomson and Bannister, 2003). Previous studies have consistently found gamma band power increases between rest and task, with no clear upper bound for frequency (Miller *et al.*, 2014). An absolute reduction of processing load in a given region should therefore manifest as a weakening of its gamma response, as the latter reflects the output of its local operations.

We presented participants with images of human faces filtered to contain either LSF only, HSF only, or both spatial frequencies (broadband). Images were either intact, i.e. containing redundancies across spatial frequency ranges, or phase-scrambled to disrupt spatial scale redundancy. Using this design, we test several predictions regarding the contribution of LSF driven feedback to visual recognition. First, if HSF processing is more efficient, i.e., recruits less computational resources, when HSF content is spatially aligned with LSF, the MEG markers of HSF processing should reduce in response to intact compared to scrambled broadband images, as found in our previous EEG study. If the early visual cortex is the primary site of such integration, reductions in HSF contribution should be found in this region. Importantly, if the decrease in HSF processing markers is due to an absolute reduction of processing load in intact, SF-redundant, viewing conditions rather than a mere shift of SF dominance, we should observe a decrease in gamma power as a proxy of reduced local processing load.

Further, if LSF feedback guides HSF processing based on an initial assessment of coarse image structure and semantic content information, it likely originates in an area that responds to visual stimuli with receptive fields that are both sufficiently large to cover the global shape of a stimulus and sufficiently abstract to represent complex objects. Ventral temporal cortex and frontal cortex (Bar, 2004; Kietzmann *et al.*, 2019) fit this profile. Ventral temporal cortical areas are involved in visual object recognition, feature large receptive fields (Sayres *et al.*, 2009; Caspers *et al.*, 2013; Kay *et al.*, 2013) and show spatial frequency sensitivity dynamics consistent with a coarse-to-fine integration of spatial scales (e.g., Goffaux et al., 2010; Musel et al., 2014). Orbitofrontal cortex (OFC) shows a bimodal spatial frequency tuning function (Fintzi and Mahon, 2014) and has also been suggested to be the source of spatial frequency specific feedback to early visual cortex (Bar *et al.*, 2006; Kauffmann *et al.*, 2015). Therefore, we expect to observe increased activity in the alpha/beta bands linked to feedback signalling, over either the inferior temporal or frontal cortex (Klopp *et al.*, 1999). Moreover, alpha/beta band feedback signals should correlate with the reduction of local processing markers in the gamma range during the processing of SF-redundant information (intact images). If instead non-semantic stimulus properties such as the alignment of edges and contours are driving the processing of redundant SF information, we expect to find such correlations with regions in the lateral occipital complex (LOC) which has been shown to be involved in resolving contours, even in the absence of semantically meaningful stimuli (Park *et al.*, 2011; Shpaner *et al.*, 2013). Systematically testing those hypotheses allows us to showcase the potential role of cross-SF-redundancy based feedback as a major influence on efficient visual computation.

## Methods

### Participants

21 participants (9 female, 12 male, mean age 28.3 SD 5.13) took part in the study in exchange for monetary compensation. If needed, corrective lenses for myopia or hyperopia were attached to the MEG helmet in front of the participants’ eyes. All participants provided written informed consent and were fully informed about the purpose of the experiment after participation. The study was approved by the Scientific Ethics Committee for the Midtjylland Region, Denmark. One participant was excluded due to excessive noise in the MEG data and one participant was excluded for failing to reach sufficient task performance.

### Stimuli

Stimuli were 47 images each of human faces, houses and their phase-scrambled versions. Houses were selected from the Pasadena houses database (Perona, 2000). We chose the 47 house images that best matched the orientation spectrum of our face stimuli. House trials were not analyzed in the current study. Faces and houses were cropped to the same oval aperture and pasted onto a uniform gray background. All image luminance histograms were matched to the same reference histogram (the average of all images) using the Shine toolbox (Willenbockel *et al.*, 2010) in Matlab. To avoid filter artifacts due to the uniform gray background, the background was replaced with a Fourier phase-scrambled version of the image by first scrambling the whole image and then re-pasting the non-scrambled aperture. Scrambled images were produced in the same way, except that the last step of re-pasting the intact aperture was skipped. To ensure that the scrambled background portrayed the low-level properties of the aperture rather than a mixture of the aperture and the original gray background the process of scrambling and re-pasting the aperture was repeated 1000 times per image thus gradually reducing the influence of the original background on the spatial frequency distribution of the scrambled background. To restrict image content to LSF or HSF, we used 7^th^ order Butterworth filters with filter cutoffs determined such that the area under the curve of the amplitude spectrum would be as similar as possible below and above the cutoff to ensure similar contrast contribution of the respective ranges to the broadband images. To avoid very low frequencies accounting for most of the contrast in the LSF images, we excluded the first 5 cycles per image (cpi) prior to determining the cutoff. This resulted in LSF passbands of > 5 and < 7cpi and HSF cutoffs of >7cpi. Finally, we masked out the phase-scrambled background with a cosine tapered mid-gray background (see Figure 1). Broadband images were constructed as the non-normalized addition of LSF and HSF images. Consequently, broadband images had higher contrast than either LSF or HSF images and importantly, because LSF and HSF images contained the same overall contrast (RMS contrast ~0.02), they contributed equally to the contrast of broadband images (RMS contrast ~0.028).

**Figure 1:**
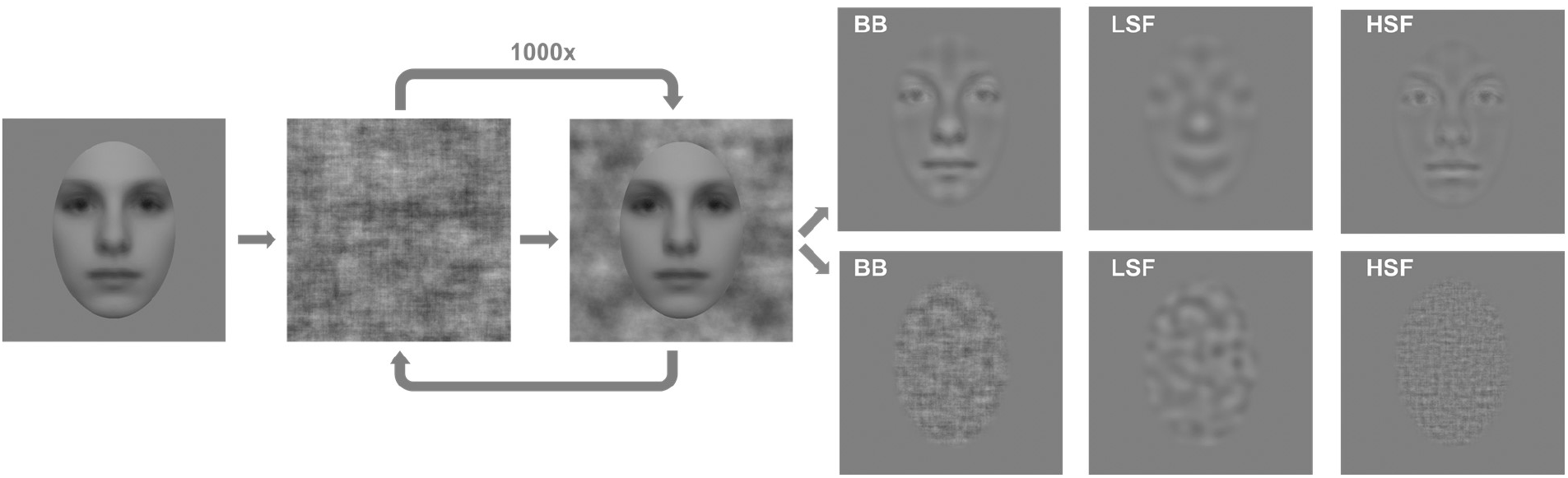
Note that example images shown here are not the original study stimuli. In order to obscure face identity for publication, the images shown here are an average of all used faces and therefore strongly biased toward low spatial frequency. The purely scrambled images reflect the true spatial frequency spectrum of the original stimuli. Stimuli were produced from full spectrum, grayscale photographs of human faces by iteratively phase scrambling the image and re-pasting the face to avoid influences of the uniform gray background on the image’s Fourier spectrum. LSF and HSF images were produced by multiplying the Fourier spectra of the iteratively scrambled images with low pass and high pass filters, respectively. The Fourier spectra of broadband (BB) images were multiplied with the sum of LSF and HSF filters without contrast normalization. In a final step, the scrambled background was replaced with gray, with the edge between face and background fading to gray over 20 pixels.

### Experimental procedure

Images were projected centrally onto a screen at 84cm viewing distance from the participant by a ProPixx projector (VPixx Technologies, Canada) in a dimly lit MEG recording chamber. The oval aperture of each image covered 22cm in height and 15cm in width resulting in a visual angle of 15° vertically and 10° horizontally. Given the degrees of visual angle covered by our stimuli, filter cut-offs translated to .46 × .7 cycles per degree.

Each participant completed 18 localizer trials and 9 experimental blocks. Every experimental block was preceded by two localizer trials, in which a sequence of images of one category (faces, houses or scrambled) was shown to the participant at a temporal frequency of 8.5 Hz (alternating image/blank screen with a 50% duty cycle). Images were band pass filtered in 18 logarithmically spaced steps covering the full range from the LSF low cutoff to the HSF high cutoff. Each sequence lasted for 18 seconds with increasing or decreasing spatial frequency (preceded by a 1sec ramp-up and followed by a 1sec ramp-down period). Participants were instructed to maintain fixation and passively view the sequences.

Experimental blocks consisted of 100 trials. Each trial started with a fixation dot on a uniformly gray screen that lasted for a jittered interval of 1.5-2.5 seconds. After that, an image of a face, a house or a phase-scrambled version of these was presented for 250 ms. Each image was filtered to contain either high, low or broadband spatial frequencies using the above-described Butterworth filters. Image presentation was followed by a blank period of 1.5 seconds. Participants were instructed to detect interspersed images showing a dog (could be in any of the three spatial frequency conditions) and report detection via a button press with their right hand. To assure participants’ attention each block included 10% target trials and participants were given short breaks between every block as well as a longer break after every three blocks.

### Electroretinogram

To provide a direct measure of retinal activity during visual stimulation, we recorded the electroretinogram (ERG) from both eyes using a corneal silver/nylon electrode (DTL Plus Electrode, Diagnosys, USA). The electrode was placed on the participant’s cornea such that it rested on the lower eyelid. Presumably due to the low contrast at high overall luminance in our stimulation paradigm, we found no discernable ERG response. Data from this recording is therefore not shown here.

### MEG acquisition and pre-processing

MEG data were acquired in a magnetically and sound shielded room from 204 planar gradiometers and 102 magnetometers (Elekta Neuromag TRIUX) at Aarhus University at a sampling rate of 2000 Hz. Eye movements were tracked via the bilateral horizontal and the right vertical EOG for later use in MEG artifact correction.

Prior to the MEG recording the position of 4 HID coils and the participant’s fiducial points as well as their individual head shape were digitized using the Polhemus Fastrak® digitizer (Colchester, VT, USA). Participants were sitting in upright position throughout the recording.

Data were offline notch-filtered to remove line noise and down-sampled to 500 Hz using MNE python (Gramfort *et al.*, 2013, 2014). Stimulus triggers were corrected to the signal of a simultaneously recorded light sensitive diode to account for delays between stimulus trigger and the first stimulus frame. Data were segmented to trials reaching from −1500 to 1500 ms around stimulus onset and visually inspected for squid jumps and obvious ocular or muscle artifacts. We excluded bad channels, artifactual trials as well as target trials from subsequent analyses.

### MRI acquisition and pre-processing

T1 weighted MRIs with an isotropic voxel size of 1 mm^3^ and a reconstructed matrix size of 256 × 256 of 15 participants were acquired via a 3 T Siemens Skyra scanner at CFIN, Aarhus University, Denmark. The remaining 4 participants provided an equivalent recent T1 weighted MRI recorded for a different study. MRIs were converted to nifty and aligned to AC/PC space using SPM 12 (Penny *et al.*, 2011).

### Sensor space analysis

Preprocessed trials were low-pass filtered by a 6^th^ order Butterworth zero phase filter with a cutoff frequency of 80 Hz, down-sampled to 250 Hz and reduced to 700 ms segments ranging from −100 ms to 600 ms around stimulus onset. Missing channels were interpolated using spherical spline interpolation of neighboring channels and planar gradients over both orientations were combined into a single magnitude measure. Data were baseline corrected to the 100 ms before stimulus onset and temporally smoothed to the moving average over three timepoints (12 ms). Regularized Linear Discriminant Analysis (LDA) using MVPA light (Blankertz *et al.*, 2011) integrated with fieldtrip (Oostenveld *et al.*, 2011) was performed on a time-point–by-time-point basis (King and Dehaene, 2014) to classify between LSF and HSF scrambled image trials. We used 5 fold cross-validation and shrinkage regularization as (1 − *λ*) * *Sw* + *λ* * *ν* * *I* where *λ* is determined by the Ledoit-Wolf estimate (Ledoit and Wolf, 2004), Sw is the scatter matrix, I the identity matrix, P is the number of features and *ν* is the trace (i.e. the sum of eigenvalues) of Sw divided by P. The trained classifiers were then evaluated on their generalization performance to broadband intact and scrambled image trials. Since the classifiers have only learned to categorize trial data as belonging to either LSF or HSF trials they will necessarily be incorrect on all broadband trials because broadband images have been specifically designed to contain equal contrast in both low and high spatial frequencies. We therefore interpret the classifier’s decision to assign either an LSF or an HSF label to a broadband datapoint as reflecting that MEG topography at the datapoint in question is more similar to one evoked by LSF or HSF stimulation, respectively, i.e., as indicating spatial frequency dominance (Petras *et al.*, 2019).Statistical significance of classification results was determined using cluster based permutation testing with alpha set to 0.01 and performing 1,000 randomizations (for details see Maris and Oostenveld, 2007).

### MEG source reconstruction

Three head position indicator (HPI) coils were used to determine the participants’ head position relative to the MEG sensors during recording. MEG data were aligned to MRI space in a three-step procedure: first the fiducial points provided by the Polhemus Fastrak® digitizer were used to calculate an initial transformation matrix and second the digitized head shape was manually aligned to a surface reconstruction of the individual’s MRI. Finally, an iterative closest point matching procedure was performed. All transformation matrices were inverted, combined into a single transformation-matrix, and applied to the MEG gradiometer descriptions separately for each run. For each participant, a three compartment boundary element head model (brain, skull, scalp) based on the individual’s MRI was created using the OpenMEEG implementation in fieldtrip (Kybic *et al.*, 2005; Gramfort *et al.*, 2010; Oostenveld *et al.*, 2011). For each participant and run, leadfields for ~42,000 positions based on a source model template warped to the individual participants’ brain were created using standard pipelines in fieldtrip. We constructed common spatial filters using a linear constrained minimum variance (LCMV) beamformer where data of all conditions was used to compute the covariance matrix. We then applied those spatial filters to reconstruct virtual channels for each grid point from the experimental trials, epoched to −500 ms-1500 ms around stimulus onset.

### ROI analysis

MEG source reconstruction has repeatedly proven to produce reliable results at spatial resolutions comparable to the one used here (Hillebrand and Barnes, 2011; Belardinelli *et al.*, 2012; Dalal *et al.*, 2013). However, errors in the alignment of MEG sensors to the participants’ MRI can cause considerable spatial inaccuracies (Hillebrand and Barnes, 2003, 2011; Dalal *et al.*, 2014). Thus, while defining regions of interest (ROIs) based on anatomical atlases is convenient, it may not provide the most responsive regions in each subject, due to (1) potential misalignment of the MEG sensors with the MRI and (2) inter-individual differences in functional maps. Here, we produced functionally-defined regions of interest for each participant using independent localizer data. Our localizer trials consisted of images flickering at a fixed frequency of 8.5 Hz with a 50% duty cycle, providing a periodic stimulation. Since the brains’ response to periodically changing stimulation is expected to show the same periodicity as the stimulus (Adrian, 1944; see Norcia *et al.*, 2015 for review), we used a single frequency dynamic imaging of coherent sources beamformer (Groß *et al.*, 2001) to localize the grid points with the highest selective response to faces. An underlying assumption of the beamformer is that sources of interest are temporally independent. Because the flicker stimulation results in very a high SNR signal that leads to highly correlated near zero-lag bilateral activation in visual cortices, this assumption is violated, potentially causing inaccurate localization. To nonetheless yield a reliable result from the beamformer, we scanned the brain volume using pairs of left/right hemisphere symmetric dipoles as the source model, a solution suggested by Popov et al. (2018). We then restricted analysis to four anatomically defined rough ROIs (calcarine fissure, fusiform gyrus, lateral occipital cortex and frontal cortex) and calculated the coherence between sources during face-image trials and the signal of a light sensitive photodiode during those same trials, normalized by the coherence between sources and diode during scrambled image trials. We used k-means clustering with 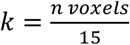 on the Euclidean distance between the 3 position coordinates and the coherence values (z-scored to ensure similar scaling of all dimensions) to parcellate sources into 4 dimensional clusters (3 position + 1 coherence dimension). To ensure matching sizes across ROIs and participants, we selected the 20 voxels with the highest coherence score from the cluster with the maximum summed coherence for further analysis. Single trial ROI virtual sensor responses were obtained by multiplying each trial’s data with the precomputed common spatial filters.

### Source space classification

We used LDA (see sensor space classification for details) to classify spatial frequency as well as image type from each ROI. The virtual channel time series between −200 ms – 700 ms around stimulus onset, derived via LCMV beamforming, served as features for the classifier. Statistical analysis of source space classification followed the same routine as described in the sensor space analysis.

### Time frequency analysis

To isolate response frequency bands of interest, we first removed the evoked response from the MEG signals by subtracting the average ERF from each trial. Then, we calculated the time frequency representation of low and high temporal frequencies separately. For low temporal frequencies we extracted wavelets linearly spaced between 3-6 cycles for temporal frequencies ranging from 2-30 Hz in steps of 1 Hz (time window −0.2-0.8 sec in steps of 0.01 sec). For the high temporal frequencies, we used discrete prolate spheroidal sequences (DPSS) multitapers (15 cycles of data at 0.2*frequency of interest smoothing) for frequencies ranging between 35 and 200 Hz in steps of 5 Hz. All power values were log transformed prior to any other analyses. To determine individual gamma frequency, we compared high frequency power between baseline (−200 to 0 ms) and the bulk of the evoked response (100 ms to 400 ms after stimulus onset) in the broadband source reconstructed ROIs using cluster-based permutation testing with dependent samples t-test on the trial-level over all conditions. Individual gamma was defined as the center frequency of the cluster with the maximum sum of t-values +/− 10 Hz. Participants where no significant cluster was found (3 out of 19 for calcarine ROI) were assigned the average of all other participants (Fusiform ROI: 86.81 Hz Lateral Occipital ROI: 99.01 Frontal ROI: 83.90 calcarine ROI: 90.64). Alpha/beta ranges were derived from previous literature (Michalareas *et al.*, 2016) and set to 8-12 Hz for alpha and 15-25 Hz for beta. We calculated average power values for all frequency bands of interest by taking the average of the (log transformed) power values and baseline correcting to the −200 to −100 ms baseline window for each ROI and condition separately.

### Gamma power comparison

We went on to test our hypothesis of reduced gamma power in response to broadband images in the intact condition in early visual cortex. First, we compared if gamma reduction in the broadband condition was stronger for the scrambled compared to the face condition. Then, we compared gamma power in the broadband versus the HSF condition separately for the intact and scrambled condition in early visual cortex (separately for each time point). All tests were corrected for multiple comparisons using cluster based permutation testing.

### Cross frequency coupling

We investigated coupling between the high-level ROIs (Fusiform, lateral occipital and frontal) and the calcarine ROI using both power-power and phase-power measures. For each ROI, we found the most representative timecourse by retaining the coefficients of the principal component that accounted for the most variance of the virtual channels within the ROI (Colclough *et al.*, 2015). We filtered this timecourse to the frequency bands of interest (individual gamma, alpha and beta) using separate low-pass and high-pass Hamming windowed one-pass zero phase-shift FIR filters with a maximum passband deviation of 0.22%, and 53dB stopband attenuation. To obtain phase estimates for the lower frequency components we chose a filter bandwidth of 4 Hz and 10 Hz respectively (α_fcenter_ 10 Hz±2 Hz; β_fcenter_ 20 Hz±5 Hz). Amplitude modulated signals feature a Fourier spectrum with a peak at the carrier frequency (i.e. the modulated frequency, here individual participants’ gamma band peak) as well as sidepeaks at the modulated frequency ± the modulation frequency (Berman *et al.*, 2012). To ensure spectral sidepeaks produced by the modulation frequency (f_mod_, α or β respectively) are included in the estimation of the higher frequency γ-power, we chose filter bandwidth to be twice the modulation frequency. As a result, the passband was defined as the center frequency of the individual participants’ γ±f_mod_ (Aru *et al.*, 2015). I.e., for a participant with a γ-peak at 70 Hz in the early visual cortex ROI, the filter for the estimation of alpha-gamma coupling has a passband of 70 Hz±f_center_(alpha) (i.e. 10 Hz) or 60 Hz to 80Hz. For that same participant, the filter for the calculation of beta-gamma coupling has a passband of 70 Hz±f_center_(beta) (i.e. 20 Hz) or 50 Hz to 90 Hz. We then calculated the Hilbert transform of the filtered time series to extract power and phase angle over time. To account for potential filter edge artifacts, we restricted further analysis to the time interval between −0.2 and 0.7 seconds around stimulus onset. For power-power coupling (PPC), we extracted the power time series for each ROI, frequency band of interest, and condition. We then correlated the γ power time series from the calcarine ROI with the α and β time series from all other ROIs using Spearman correlations. For phase-amplitude coupling (PAC), we extracted γ-power time series from the calcarine ROI and α and β phase time series for all other ROIs. We calculated phase amplitude coupling as the length of the average vector of γ-power and lower frequency band phase in polar space. Because our hypothesis was that EVC γ-power is influenced by α or β power or phase in high-level ROIs, and transmission velocity in the brain is finite, we expected correlations between ROI timeseries to be maximally different from baseline when the EVC timeseries was lagged with respect to the high-level ROI timeseries. We therefore calculated PAC and PPC for 5 different lags ranging from −40 ms (high level ROI leading) to +40 ms (EVC leading) in steps of 10 ms. To correct for potential confounds due to outlier power fluctuations or violations of the von Mises distributions (in the case of PAC) we standardized individual subject PAC and PPC values using the mean and standard deviation of the permutation distribution under the null hypothesis (no temporal relationship between frequency bands). To achieve this, we temporally shifted the lower frequency band power timeseries with respect to the gamma band timeseries by random offsets as suggested by Cohen (2014). We performed 1000 iterations per timepoint and lag. On the group level, we tested for the interaction between image type and spatial frequency condition as well as the main effect of either manipulation using cluster statistics (with dependent samples F-test (interaction) or t-test (main effect) as statistics, at an alpha of 0.05 with 1000 permutations).

## Results

We investigated the signaling dynamics between early visual cortex and inferior temporal cortex, as well as frontal cortex using MEG. Participants passively viewed images of intact and phase-scrambled images of faces with similar amplitude and orientation spectra for 250 ms each. Images were filtered to contain either low, high, or both (broadband) spatial frequencies. The filter cutoffs were chosen such that LSF and HSF components contributed equally to the overall contrast in the broadband images (see Methods). Phase-scrambled images were produced by iteratively Fourier-scrambling images and re-pasting the original face to ensure that amplitude and orientation spectra were closely matching those of the original images and not influenced by the uniform gray background (see Figure 1 and Methods).

### Sensor space spatial frequency dominance patterns are modulated by image coherence

Tracking the dynamics of spatial frequency integration in naturalistic broadband images is complicated by the need to disentangle the contribution of each frequency band without interfering with the integration itself. We have previously demonstrated that multivariate classifiers trained to distinguish between patterns evoked by either frequency band in isolation can be used to track spatial frequency processing dominance during broadband stimulation (Petras *et al.*, 2019). Here, we replicate these findings in a different group of participants, using MEG instead of EEG.

We trained linear discriminant analysis (LDA) classifiers to perform a time-point-by-time-point discrimination of LSF and HSF phase-scrambled image trials to concentrate on spatial frequency encoding per se without any potential influence of image semantic content on classification. The distribution of magnetic field changes measured by a set of 204 planar gradiometers served as features for the classification (see Figure 2 for topographies and event-related fields).

**Figure 2:**
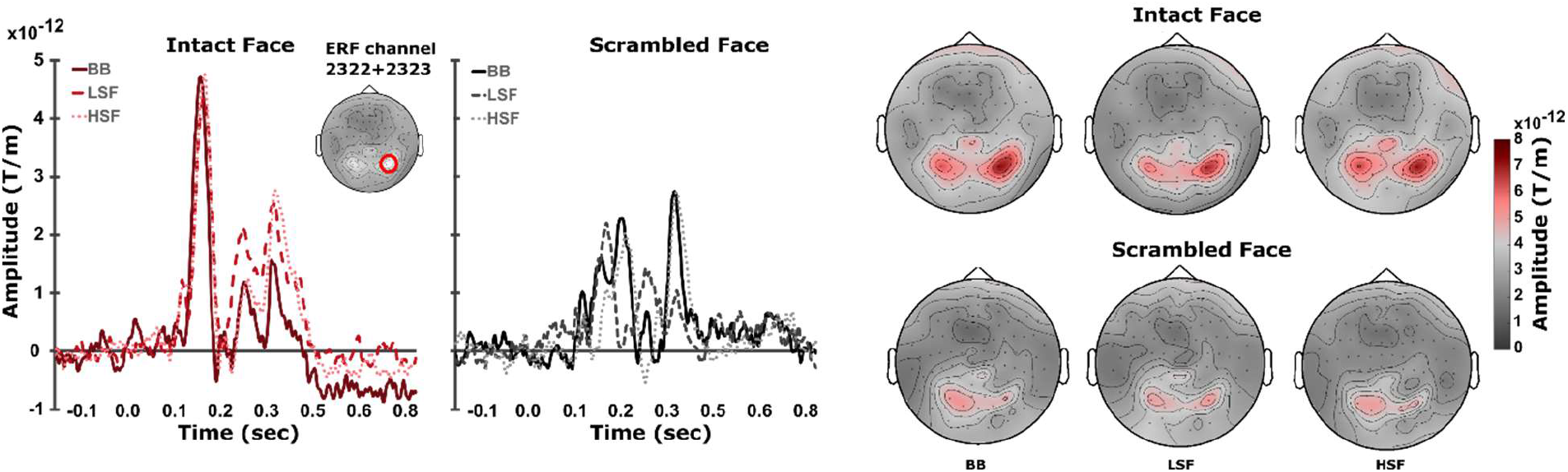
Sensor space baseline corrected event-related fields (ERFs) and topographies for all conditions. Left: ERFs at combined gradiometers 2322 + 2323 (red circle). Right: Topographies at the latency of the first response peak (150 ms ±15 ms after stimulus onset).

Classification performance was significantly above chance as revealed by a cluster-based permutation test [p < 0.001, cluster statistic = 26327 (maximum sum of cluster t-values)] and first exceeded chance levels around 90 ms after stimulus onset (see Figure 3a+b). Performance remained above chance until about 450 ms after stimulus onset (i.e. 200 ms after stimulus offset). As expected, classification performance was highest when training data and cross-validation data were sampled from the same latencies in trial-time as seen along the diagonal in Figure 3a. However, there was considerable cross-time generalization (off-diagonal decoding), indicating a sustained spatial frequency specific pattern response.

**Figure 3:**
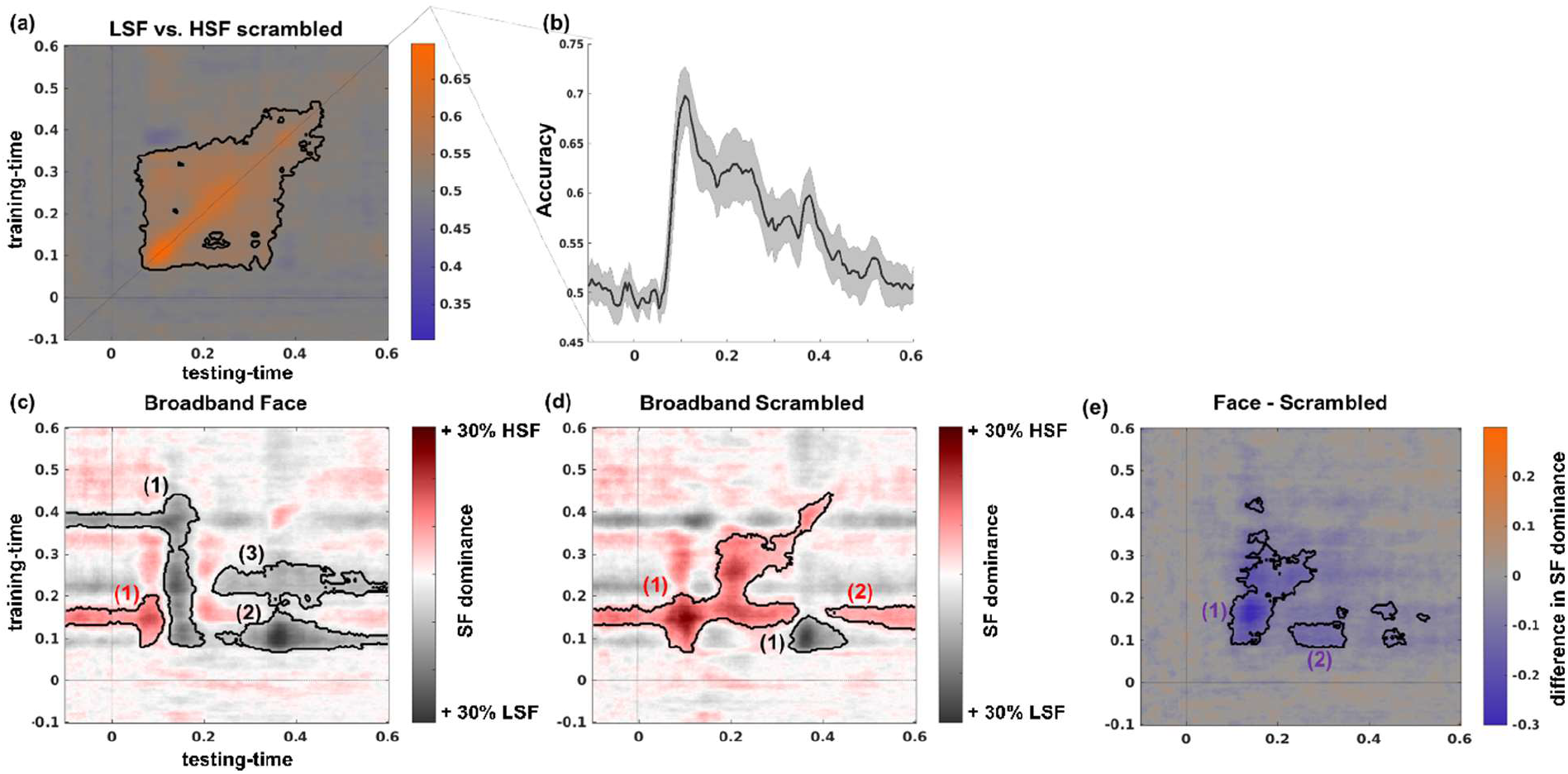
Sensor space spatial frequency classification and generalization. (a) Time point-by-time point spatial frequency decoding. (b) Classification performance when training time and testing time are the same (diagonal of (a)). (c) & (d) Generalization performance of classifiers trained in (a) when evaluated on broadband trials. SF dominance indicates the proportion of LSF or HSF decisions the classifier made. (c) Generalization performance for broadband intact face trials (d) Generalization performance for broadband scrambled trials. (e) Difference of classifier generalization between intact face and scrambled trials. Purple colors indicate stronger LSF dominance for intact face trials and/or stronger HSF dominance for scrambled trials. Dark contours delineate statistically significant clusters. Cluster indices correspond to the statistics reported below.

We evaluated how the classifiers pre-trained to differentiate between LSF and HSF scrambled image trials performed on broadband trials (Figure 3). In the latter case the classifiers response reflects which of the two spatial frequency ranges is dominantly represented in the scalp pattern evoked by the broadband trials. For both broadband intact and scrambled trial data, the classifiers responses significantly deviated from chance towards LSF, as well as towards HSF in separate clusters (see Figure 3c–d and Table 1).

**Table 1:**
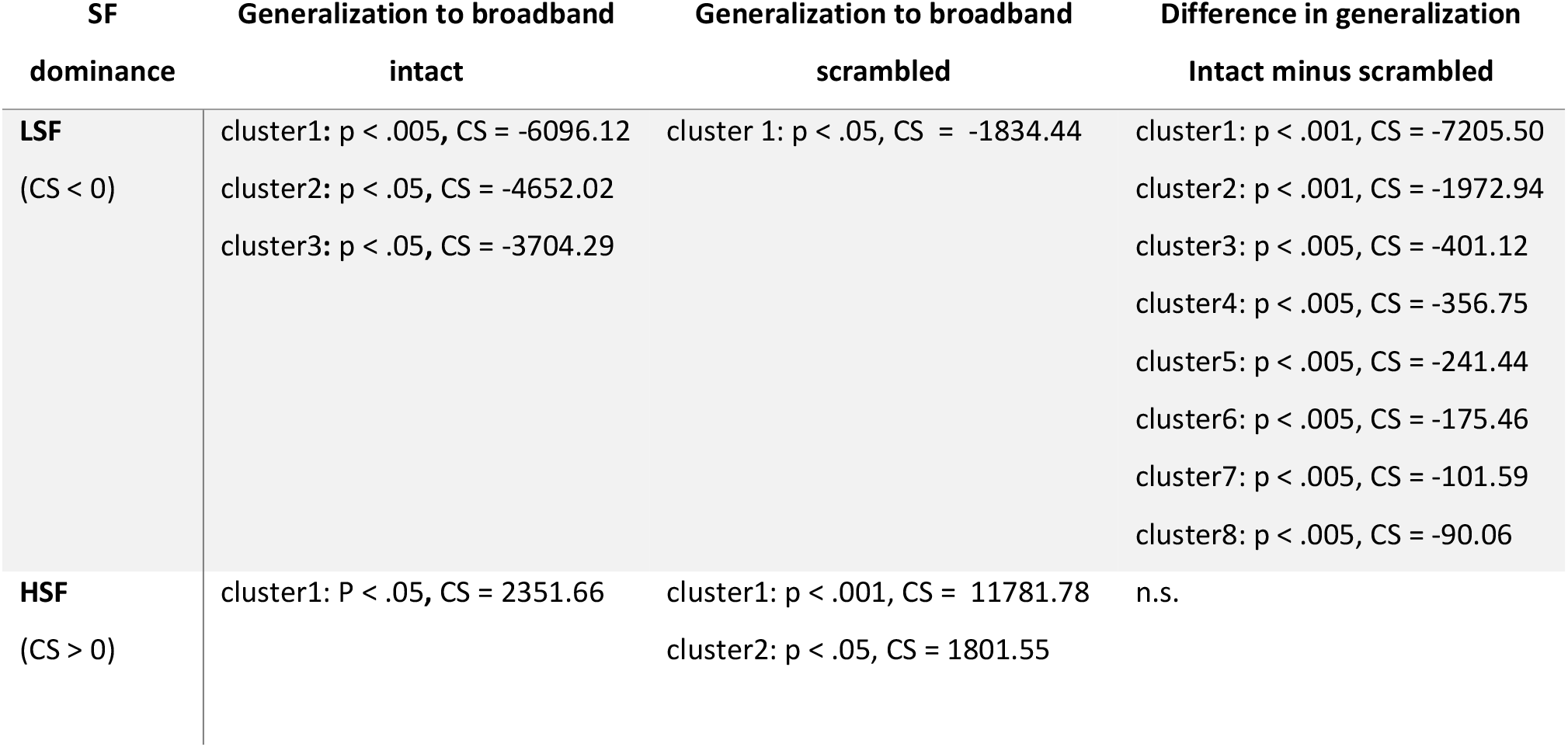
Sensor space LSF vs HSF trial classification results. LDA classifiers were trained on scrambled image trials to differentiate between LSF and HSF image trials. CS = cluster statistic (maximum sum of cluster t-values). Classifier’s generalization bias reflects SF representational dominance with negative values for CS indicating LSF dominance. Cluster numbers are in descending order of maximum cluster statistic.

Importantly, intact and scrambled image processing differ significantly in their spatial frequency dominance with intact faces showing lower HSF dominance (i.e., greater LSF dominance) than those evoked by scrambled images (see Figure 3e). The reduced HSF dominance in response to intact, compared to scrambled images suggests that when LSF aligns in space with HSF, LSF information guides HSF processing, thus alleviating computational resources. These results replicate our previous findings on spatial frequency dominance obtained from EEG recordings. Leveraging the superior spatial specificity that MEG data provides, we next reconstructed the cortical sources underlying the observed effects.

### Functional localizers for individual ROIs

We hypothesized that LSF driven feedback modulates early visual cortex processing when SF are aligned in the stimulus, i.e., when LSF is informative towards HSF. To constrain our analysis to the relevant cortical regions according to our hypothesis, we performed source reconstructions on our data. We used functional localizer trials to restrict regions of interest (ROIs) to areas that preferentially responded to images of faces. Basing the selection of ROIs on functional localizers obtained in the same session as the experimental data is more robust against small errors in head-to-MEG helmet alignment, assuming the participants head position was stable within runs (see methods for details).

Prior to each block of the main experiment, participants watched two sequences of either face or scrambled image stimuli lasting for 20 seconds each (see methods for details). In order to maximize efficiency and obtain high SNR estimates from limited recording time, we used a periodic visual stimulation technique (Norcia *et al.*, 2015) where images were flickered at a steady rate of 8.5 Hz (17 Hz with a 50% duty cycle). Such periodic stimulation produces the steady state visual evoked response (Adrian, 1944; Peli, McCormack and Sokol, 1988; Strasburger, Scheidler and Rentschler, 1988), a strong oscillatory potential at the stimulation frequency and its harmonics (see Figure 4a for sensor space representation).

**Figure 4:**
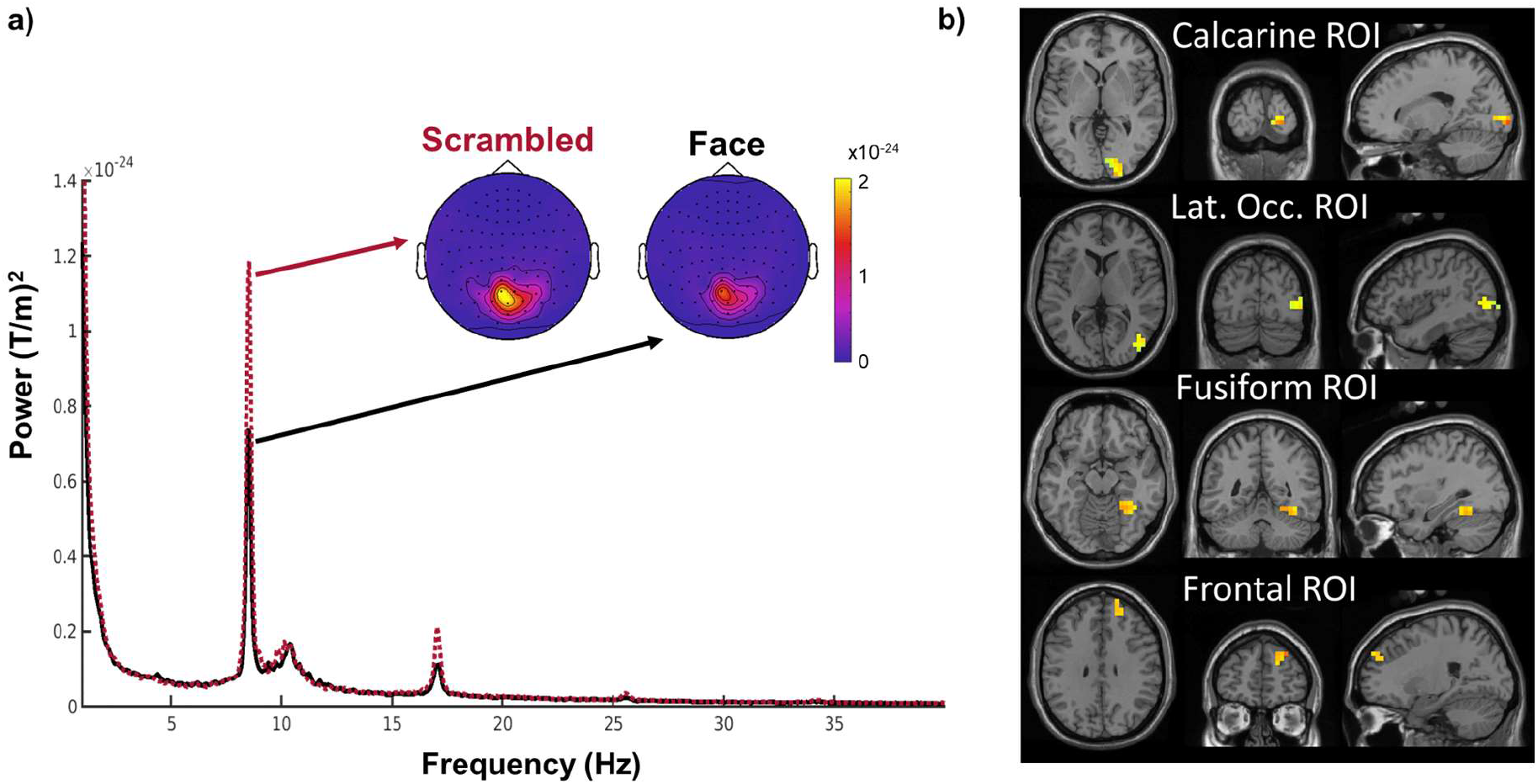
Functional localizers for individual ROIs. Figure 4: a) Flickering stimuli at a fixed frequency (here 8.5 Hz) evokes a strong response at the stimulation frequency and its harmonics averaged over all occipital channels. b) Regions of interest for an example participant. Marked are the 20 most responsive voxels per ROI. Images are in neurological convention. For all ROIs of all participants see supplementary Figure 1.

We used dynamic imaging of coherent sources (DICS, Groß *et al.*, 2001) to reconstruct the source response that was phase-locked to a simultaneously recorded photodiode signal by calculating the coherence coefficient between each grid point in source space and the photodiode signal. We then defined four rough anatomical regions of interest: the calcarine fissure, the lateral occipital cortex, the fusiform gyrus, and the frontal cortex. To constrain our analysis to regions with a robust response to face stimuli, we used k-means clustering to select the cluster of the 20 source dipoles with the highest summed coherence in response to faces for each participant within each of these coarse parcels, (see Figure 4b and Methods).

### Source space classification reveals reduced high spatial frequency dominance for intact images

To test our hypothesis that LSF driven feedback reduces the HSF related activation in early visual cortex, we applied the same classification analysis we performed on the sensor space data, on each of the source space ROIs. Classification performance significantly exceeded chance levels in all ROIs when differentiating between low and high spatial frequencies of scrambled stimuli (Figure 5 left column; also see table 2), indicating that all tested areas are sensitive to spatial frequency differences, even in the absence of meaningful shapes or semantic information.

**Table 2:**
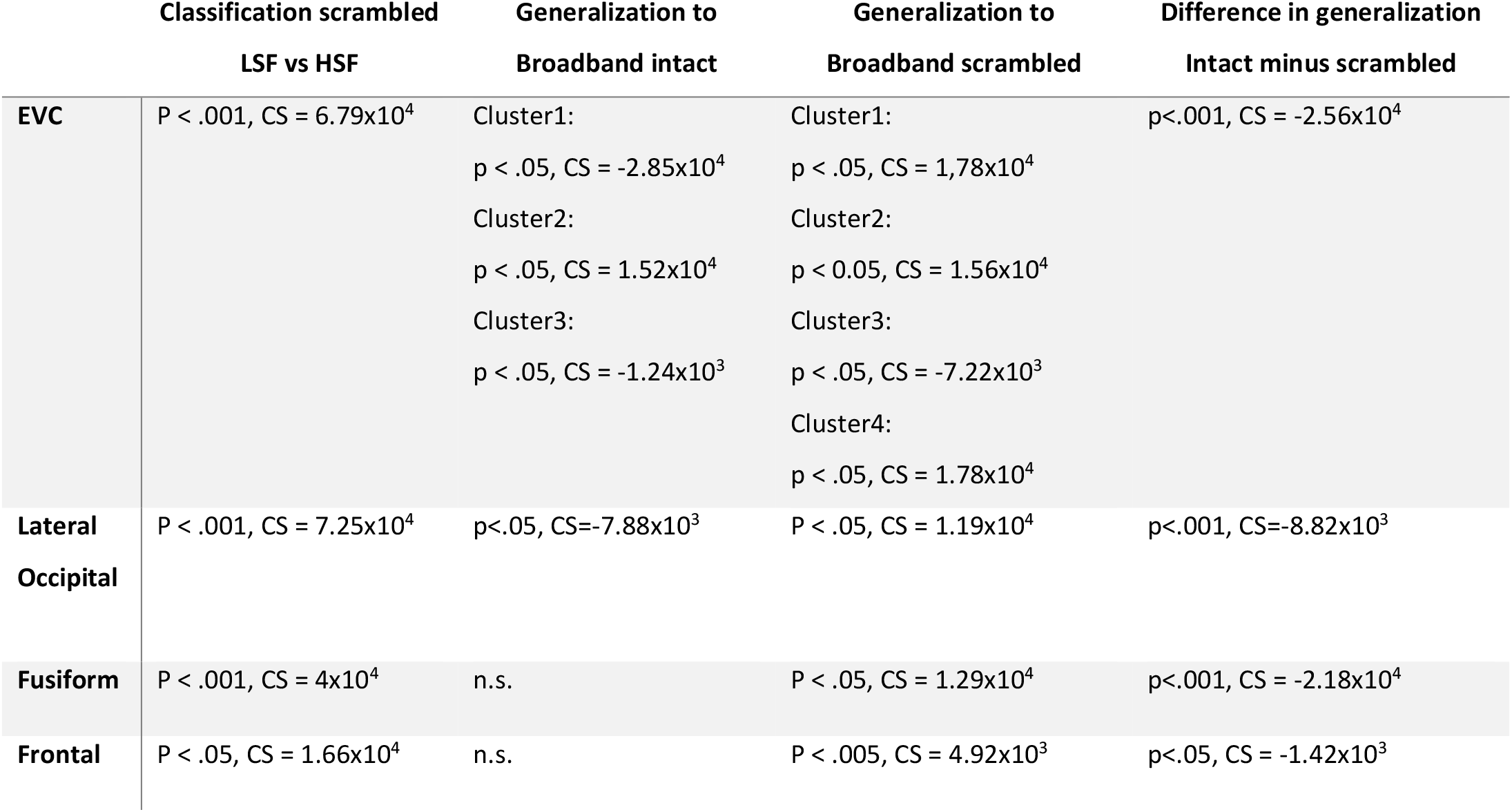
LSF vs HSF trial classification results. LDA classifiers were trained on scrambled image trials to differentiate between LSF and HSF image trials. CS = cluster statistic (maximum sum). Classifier generalization bias reflects SF representational dominance with negative values for CS indicating LSF dominance.

|  | Classification scrambled<br>LSF vs HSF | Generalization to<br>Broadband intact | Generalization to<br>Broadband scrambled | Difference in generalization<br>Intact minus scrambled |
| --- | --- | --- | --- | --- |
| **EVC** | P < .001, CS = 6.79x10<sup>4</sup> | Cluster1:<br>p < .05, CS = -2.85x10<sup>4</sup><br>Cluster2:<br>p < .05, CS = 1.52x10<sup>4</sup><br>Cluster3:<br>p < .05, CS = -1.24x10<sup>3</sup> | Cluster1:<br>p < .05, CS = 1.78x10<sup>4</sup><br>Cluster2:<br>p < 0.05, CS = 1.56x10<sup>4</sup><br>Cluster3:<br>p < .05, CS = -7.22x10<sup>3</sup><br>Cluster4:<br>p < .05, CS = 1.78x10<sup>4</sup> | p<.001, CS = -2.56x10<sup>4</sup> |
| **Lateral Occipital** | P < .001, CS = 7.25x10<sup>4</sup> | p<.05, CS=-7.88x10<sup>3</sup> | P < .05, CS = 1.19x10<sup>4</sup> | p<.001, CS=-8.82x10<sup>3</sup> |
| **Fusiform** | P < .001, CS = 4x10<sup>4</sup> | n.s. | P < .05, CS = 1.29x10<sup>4</sup> | p<.001, CS = -2.18x10<sup>4</sup> |
| **Frontal** | P < .05, CS = 1.66x10<sup>4</sup> | n.s. | P < .005, CS = 4.92x10<sup>3</sup> | p<.05, CS = -1.42x10<sup>3</sup> |

**Figure 5:**
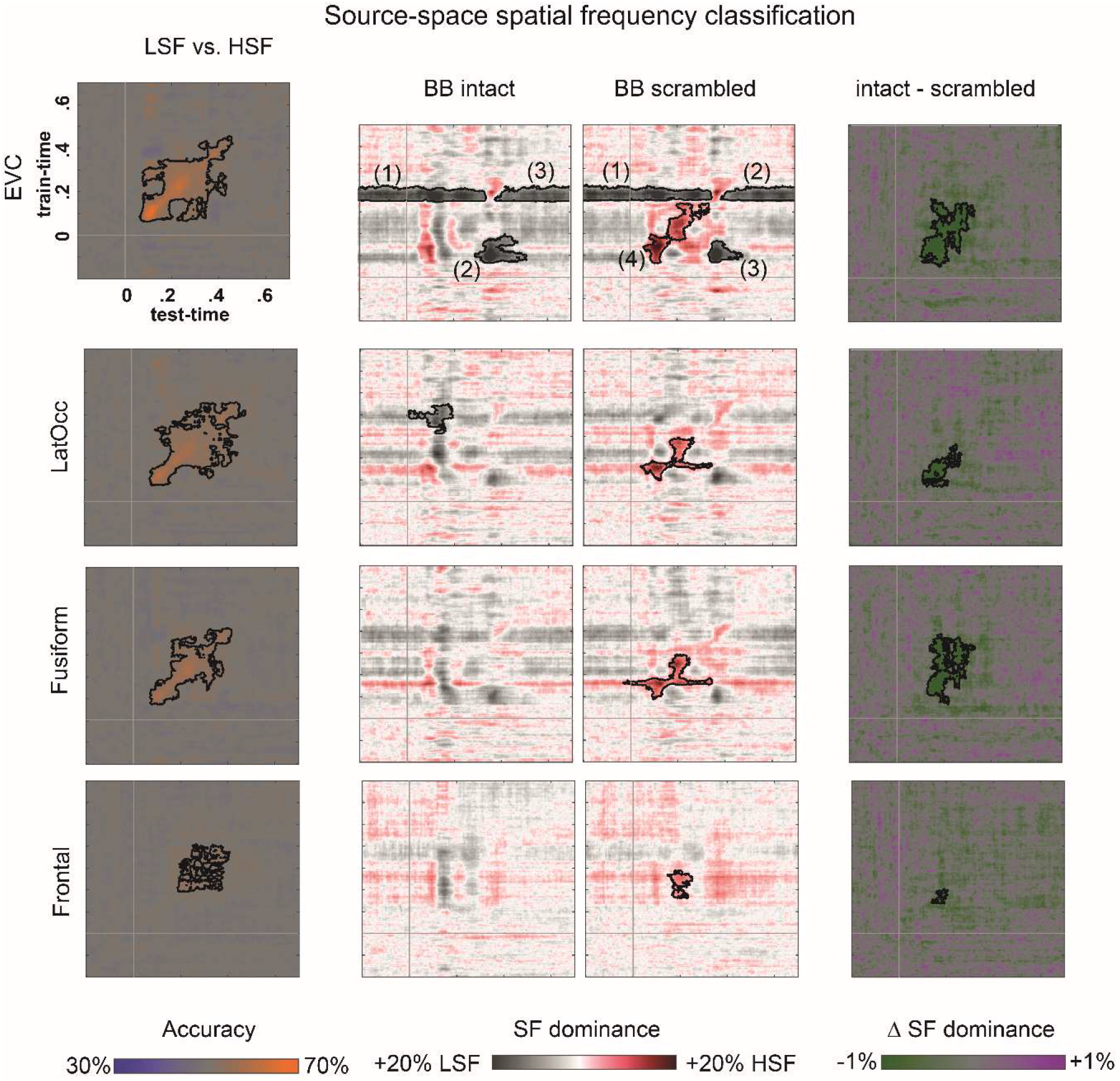
Spatial frequency classification and generalization in source-space. All four ROI contain sufficient information to differentiate between scrambled LSF and HSF trials, although accuracies and latencies of successful classification differ substantially across them. When applying the classifiers trained on LSF vs HSF scrambled image trials to broadband image trials, the early visual cortex ROI shows a similar SF dominance pattern as observed in sensor-space with stronger HSF dominance in response to scrambled, compared to intact face images. Cluster indices reflect order in ranked cluster statistic (see table 2).

Next, we investigated the contribution of individual spatial frequency bands to the activation pattern evoked by broadband images, in which scales were either aligned and therefore providing redundant information (i.e., intact images), or not (i.e., scrambled images). Coarse-to-fine guidance is only expected in the intact condition. To this aim, we applied the classifiers trained on LSF vs HSF scrambled image trials to broadband intact versus scrambled images. For intact images, early visual cortex and lateral occipital ROIs both show significant LSF dominance (Figure 5). The same classifiers applied to scrambled image trials reveal both LSF and HSF dominant periods in the early visual cortex ROI whereas only HSF dominance periods were found in the Fusiform, Lateral Occipital and the Frontal ROIs. In contrast to the more common use of classification techniques in cognitive neuroscience, where the classifier output is evaluated against known class labels, training on LSF vs. HSF trials and evaluating on broadband trials, where LSF and HSF contribute equally, confronts us with a classification problem that is ill-posed: SF dominance classification cannot be compared to a known ground truth. To verify that classification of source-reconstructed responses is feasible and generalizes across spatial frequency conditions, we constructed a generalization problem with a known ground truth by classifying between intact and scrambled images within and across SF conditions. We found classification as well as generalization performance to be significantly above chance in all investigated ROIs (see supplementary figure 2 for full results).

In all investigated ROIs, HSF dominance was reduced in intact compared to scrambled images indicating either an absolute reduction of HSF related processing, or a relative shift of processing resources away from HSF, toward LSF components. To disambiguate between those alternatives, we turn to an analysis of gamma band power in early visual cortex (see below).

### Reduced gamma response in early visual cortex when informative LSF is available

We hypothesized that under broadband intact image conditions, which allow for the LSF guidance of HSF processing, the resources dedicated to HSF processing in early visual cortex are reduced. To test this, we compared the power of the neural response in the gamma frequency range (individual participants γ-peak ±10 Hz) as a proxy of local processing for each participant across conditions (see Methods for details).

Time frequency analysis revealed an interaction between spatial frequency and image type condition [BB intact – HSF intact vs. BB scrambled – HSF scrambled, CS=220.48, p<.05, cluster corrected for multiple comparisons]. When directly comparing broadband intact and scrambled images (see Figure 6a), the early visual cortex ROI showed a decrease in gamma power in response to intact compared to scrambled images [CS=−42.9, p <.05]. This effect was not found for either spatial frequency in isolation [max CS LSF =4.7, p>0.05; max CS HSF = 2.7, p>0.05]. We interpret this finding as supporting a role of LSF information in the more efficient integration of HSF image detail in early visual cortex when LSF information contained in the broadband images was informative towards the content of HSF information (i.e. in intact, but not in scrambled images).

**Figure 6:**
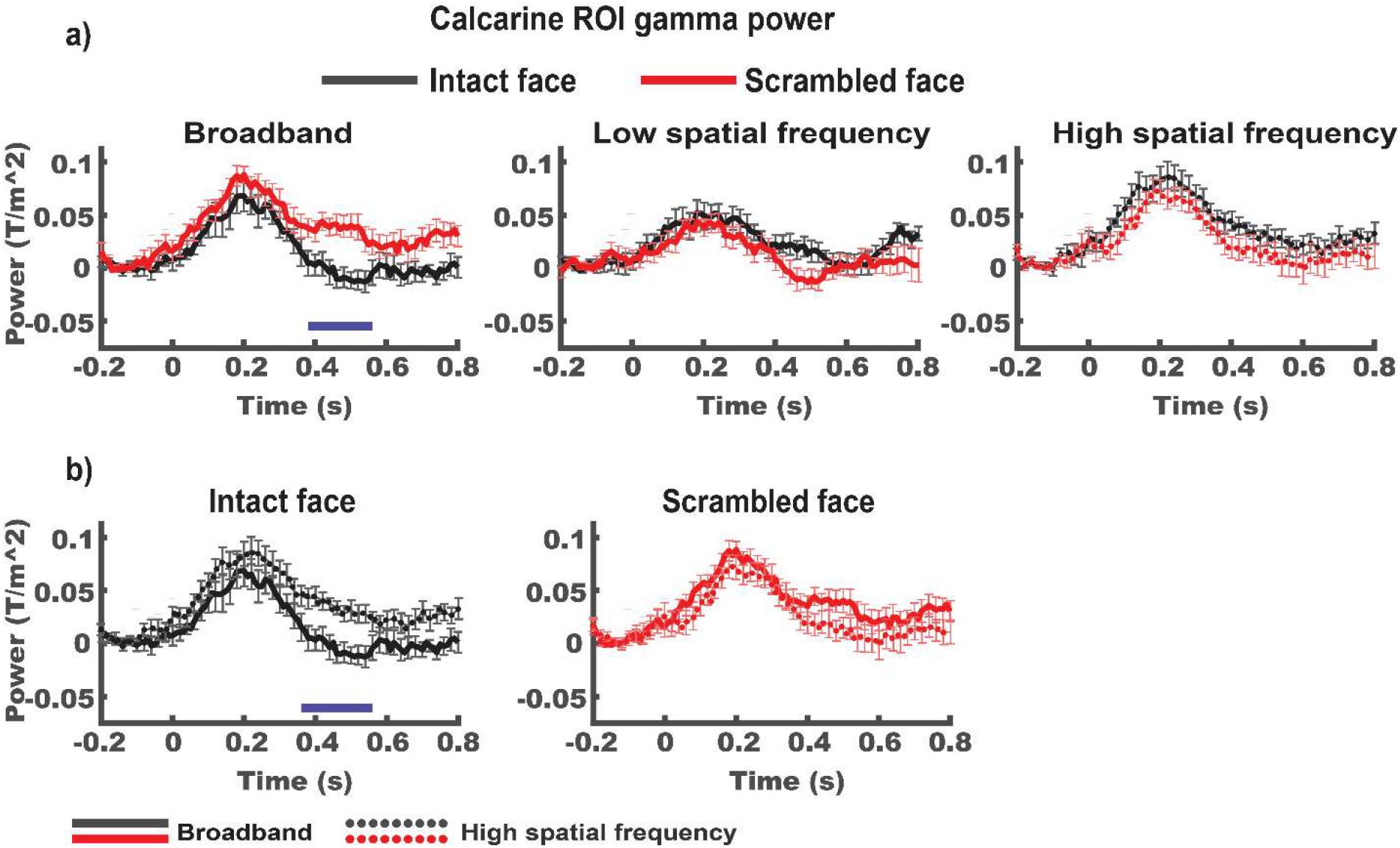
a) γ-band source power in the calcarine ROI is significantly lower in response to intact, compared to face scrambled stimuli only when images contain broadband spatial frequencies [p<0.05, the blue horizontal bar indicates the temporal extend of the corresponding cluster]. Note that responses to LSF stimuli are shown here for completeness but were not part of the planned interaction analysis. b) Direct comparison between broadband and high spatial frequency conditions for intact left panel) and scrambled (right panel) trials. Data is re-plotted from a). Note that during high spatial frequency processing, γ-power in response to intact images exceeds the maximum reached during broadband stimulation [p<0.05, the blue horizontal bar indicates the temporal extend of the corresponding cluster]. All data are baseline adjusted to the 200 ms prior to stimulus onset. Error bars indicate standard error of the mean.

Simple effects analysis of spatial frequency condition (see Figure 6b) showed lower gamma power for BB, compared to HSF trials when face images were intact [CS=−57.8, p<.05], but not when images were scrambled [CS=2.7, p>0.05]. This result is striking because broadband images had a higher overall contrast than both LSF and HSF images alone and brain responses to visual stimulation usually increase as a function of contrast (e.g., Gebodh, Vanegas and Kelly, 2017).

The finding that scale alignment in broadband images decreases the power of gamma band response in early visual cortex indicates that the presence of informative LSF content in intact broadband images reduces the lower-level computational resources recruited for the processing of HSF content. In other words, LSF-driven guidance makes the processing of HSF more efficient in early visual cortex. In the next section we investigate whether such LSF-driven guidance is carried by feedback signals from higher-level visual regions.

### Reduced gamma response in early visual cortex does not show strong correlations with alpha/beta power or phase in any of the tested ROIs

To test our hypothesis that feedback from high-level visually responsive regions is driving the more efficient processing of intact compared to scrambled images, we investigated the relationship between alpha/beta band responses, as a proxy of feedback processes, and early visual cortex gamma power. Supplementary Figure 3 provides an overview of alpha and beta power in all ROIs and conditions. Because cross-frequency coupling effects can be found in both power and phase correlations (Canolty *et al.*, 2006; Siegel, Warden and Miller, 2009; Canolty and Knight, 2010), we tested for both power-to-power and phase-to-(squared) amplitude coupling (PPC and PAC, respectively).

For the PPC, we correlated the amplitude envelopes of the early visual cortex gamma band to alpha and beta band power of all other ROIs using lagged correlations. We expected higher alpha/beta power in upstream regions (“more feedback”) to predict lower gamma power in early visual cortex (“reduced processing load”), i.e. more negative correlations for intact compared to scrambled image trials. We produced individual subjects z-scores for each timepoint-by-lag correlation (see methods for detailed description of the analysis and supplementary figure 4a for complete result). For brevity, Figure 9a shows a condensed version of the full analysis where lag between higher level alpha and beta band power and EVC gamma is zero. We found no significant correlations after correcting for multiple comparisons (see methods for detail).

**Figure 9:**
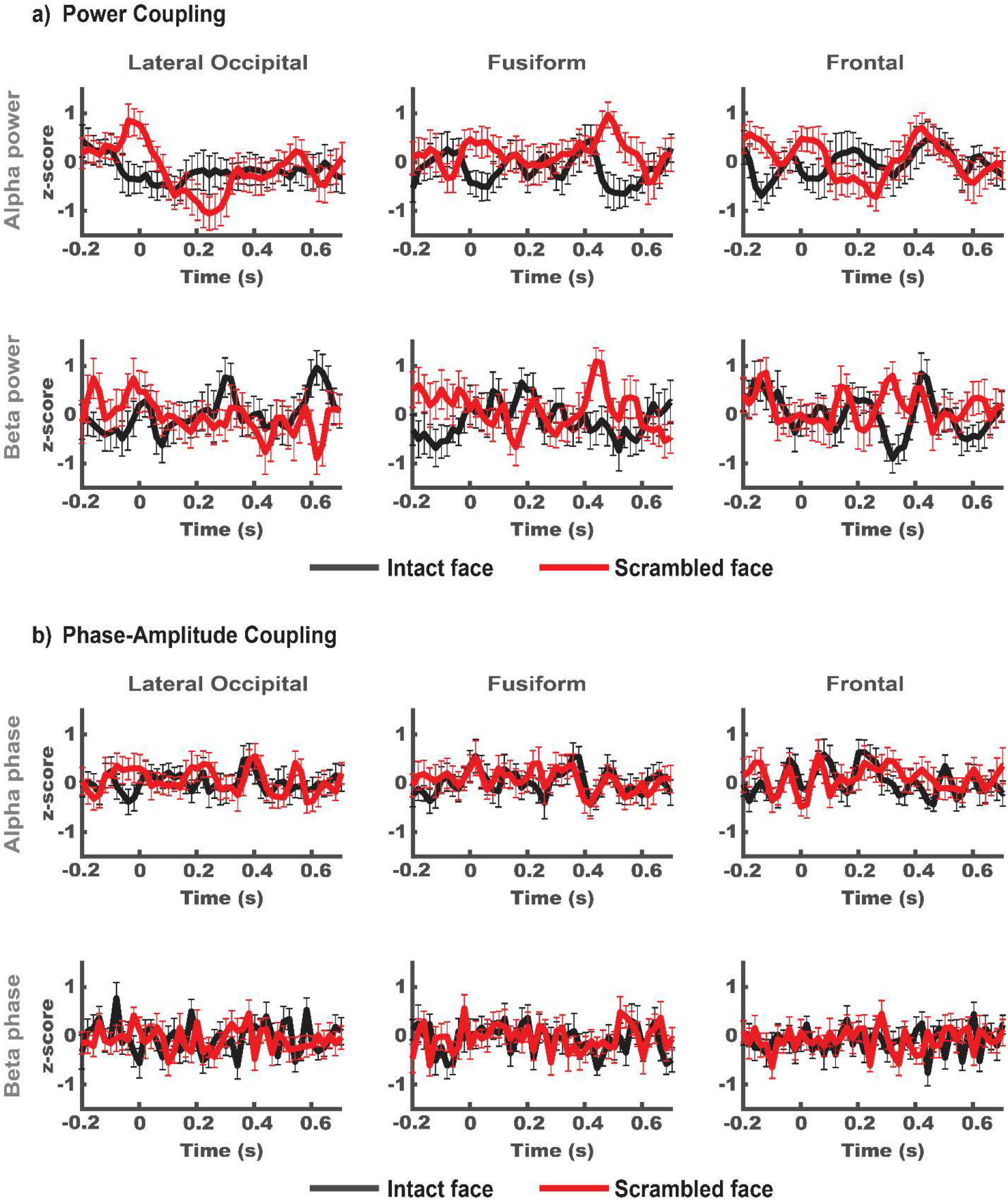
a) Standardized power-power coupling (PPC) of early visual cortex ROI γ-band and the α- and β-bands in all other ROIs. Differences between conditions do not meet our significance threshold (p<0.05) after correction for multiple comparisons. b) Standardized phase-amplitude coupling (PAC) between early visual cortex ROI γ-band and the α- and β-bands in all other ROIs. We found no significant differences between conditions. Statistical analysis was performed taking lagged correlations (±40 ms) into account. For brevity, only results for broadband image trials and 0 lag are plotted here. All other conditions can be found in the supplementary materials. Error bars indicate standard error of the mean.

For the PAC, we estimated phase amplitude coupling between instantaneous alpha and beta phase of all ROIs with gamma power of the early visual cortex ROI. We found no significant difference between estimated PAC and the permutation distribution.

To compare between conditions, we ran cluster-based permutation tests for the interaction between stimulus bandwidth (broadband - HSF) and stimulus type (intact – scrambled) as well as simple effects of spatial frequency and stimulus type on both power-to-power and phase-to-power coupling. If decreases in EVC gamma power in response to intact broadband, compared to scrambled or HSF images were driven by feedback signaling from upstream ROIs, we would expect increased PAC and/or a negative PPC relationship between upstream areas and early visual cortex. However, we found no significant difference in PAC between conditions and only a trend for differences in PPC for Fusiform-EVC between intact and scrambled broadband conditions (Figure 9a). Once we corrected for the number of tests, this trend did not meet our significance criterion of p<0.05. Altogether our results offer no support for our hypothesis that the reduction of EVC gamma power we observe for intact compared to scrambled broadband images relies on feedback from any of the ROIs we investigated (but see discussion for methodological considerations).

## Discussion

We used MEG to track the neural signatures of LSF and HSF processing in the visual responses to broadband images of intact and phase-scrambled faces. The consistent spatial alignment of contrast across spatial frequencies in natural (intact) images constitutes information redundancies insofar that LSF can be used to index the portions of HSF that are most relevant for the perceptual task at hand. Here, consistent with our previous EEG results, we showed that during the viewing of broadband face images, markers of HSF processing were reduced when images were intact, i.e., featuring spatially aligned low spatial frequency contrast, compared to when image phase (and with it spatial alignment across scales) was scrambled. We interpret our results to suggest that the presence of informative LSF information in intact broadband images indexed and facilitated the processing of their HSF content. This interpretation is supported by our finding that the reduced HSF dominance for scale-redundant images goes along with a reduction of gamma range responses in the early visual cortex. Since gamma band oscillations presumably reflect local feedforward signaling, the decrease of gamma power in early visual cortex suggests that our past EEG and current MEG classification findings reflect a genuine reduction of HSF processing load for intact, SF-redundant, viewing conditions, rather than a mere shift of SF dominance. This result corroborates a core tenet of coarse-to-fine theories of vision, namely that prior LSF indexing facilitates the subsequent encoding of HSF image content. However, the interpretation of our results requires careful consideration of potential pitfalls and alternative accounts. We here address theoretical and methodological considerations.

### Gamma effects and microsaccades

Consistent with our past EEG findings, we find the effects of image phase scrambling to be relatively late in the visual response (early visual cortex gamma power reduction for intact vs scrambled around 400 ms), although an exact temporal localization would require different tradeoffs between time and frequency resolution than the ones we chose here. The long latencies and sustained differences between conditions are in contrast with the typically shorter latencies and transient responses of layer 4 neurons typically receiving feedforward input (Nowak *et al.*, 1995). They might thus reflect EVC output after consolidation, supporting a sustained, potentially recurrent contribution to perception (Hegdé, 2008).

An alternative account for the observed changes in broadband induced gamma activity is that they do not reflect oscillatory neuronal activity at all, but are instead muscle artifacts caused by microsaccades (Yuval-Greenberg *et al.*, 2008; Melloni *et al.*, 2009). We have several reasons to believe that our findings are not the result of such saccadic artifacts: First, Yuval-Greenberg et al. and Melloni et al. demonstrated the saccadic confound on transient broadband gamma band responses recorded with EEG with a nose reference. But even with average referencing, ocular artifacts in EEG data cannot be fully discounted because volume conduction complicates reliable source reconstruction (De Munck and Van Dijk, 1991). MEG, as used in the current study, does not suffer equally from such confounds since MEG does not require a reference channel. Further, while ocular artifacts might still contaminate MEG data, their localization and removal is comparatively straightforward (Carl *et al.*, 2012).

Nevertheless, the possibility that microsaccades contribute to the observed differences in gamma band responses remains. In a study by Craddock et al. (2017), the authors show stimulus dependent changes in microsaccade frequency and amplitude using visual stimuli similar to the ones used here. Contrary to our gamma band results, they report higher rates of microsaccades for intact, compared to scrambled objects as well as for broadband and high spatial frequency compared to low spatial frequency intact images. No significant difference in saccade rates was observed between broadband and HSF images. We find *decreases* in gamma responses for intact compared to scrambled images. Consequently, even if present in our data, we would expect potential microsaccade contributions to counteract rather than facilitate the gamma band differences we found. Lastly, microsaccade rates have repeatedly been shown to rebound to above baseline peaks after an initial dip with stimulus onset. The latency of this rebound is typically reported as ~200-300 ms post stimulus onset (Schwartzman and Kranczioch, 2011) while the gamma band differences we find here are later in trial time (again, with the caveat of time-frequency-resolution tradeoffs).

### Information redundancy or basic image statistics?

We demonstrated that image scrambling influences the HSF representational dominance as well as EVC gamma power. Scrambling the phase of an image disrupts the alignment of its spatial scales while leaving other first- and second-order statistical regularities such as mean luminance, contrast, spatial frequency, and orientation spectrum unaffected (see for example Honey, Kirchner and VanRullen, 2008). It is therefore a reliable means to disrupt the integration of SF while preserving the global SF power spectrum.

In interpreting differences in the neural responses to intact and scrambled images, it is however important to note that the disruption of continuous edges introduced via phase-scrambling also alters image sparseness and might lead to processing differences due to e.g. lateral inhibition processes within early visual regions (see e.g. Gregor, Szlam and LeCun, 2011). We consider it unlikely that the reduction of power in the gamma band we found for BB intact compared to scrambled images can be fully explained by differences in image sparseness since a similar effect was not found in LSF or HSF intact compared to scrambled images that should suffer from the same sparseness confound.

A further concern is that the frequency of gamma responses increases with stimulus contrast as Hadjipapas et al. (2015) cautioned. To ensure that LSF and HSF images were fully represented in our broadband stimuli, we constructed broadband images by adding the LSF and HSF without normalization. This procedure leads to broadband images having higher RMS contrast than both LSF and HSF alone. It is therefore possible that our approach to determine individual gamma frequency based on each participant’s average response over all conditions deviated from the optimal gamma frequency for each individual condition since the higher contrast condition (i.e. broadband) may weight stronger than the lower contrast conditions. Nonetheless we found early visual cortex gamma power to be lower in response to intact broadband images compared to intact HSF images. Further, we find it unlikely a potential increase in gamma frequency in the broadband condition would have been sufficiently severe to remove the response of interest out of our chosen individual gamma range (individual gamma peak ± 10 Hz). In the above-mentioned study, the average frequency-increase the authors found, even after correcting for the dominating power increase, was 8 HZ for a contrast difference of 60%. The RMS contrasts of our stimuli are ~0.02 for narrowband and ~0.028 for broadband images (40% increase). Lastly, we find the effect of reduced gamma power in broadband compared to HSF images in response to intact images only. Gamma power responses to scrambled images, that have the same differences in RMS contrast, but contain no redundant information across spatial frequency bands, do not differ between SF conditions. We therefore maintain our interpretation that observed gamma differences are best explained by the presence or absence of cross spatial frequency information redundancies in the visual stimulation.

### LSF driven guidance of HSF processing in early visual cortex

Coarse-to-fine models of vision propose that image low spatial frequencies serve to guide HSF processing based on their initial assessment by high-level visual regions equipped with receptive fields that are sufficiently large and abstract to capture the global image structure and semantic information. In this framework, the result of this initial computation is presumably fed back to primary visual regions to better target resources for subsequent HSF processing. In the example of face perception, an efficient way to integrate different ranges of spatial frequencies would thus be to use the initial representation of image LSF to first detect the presence of a face (Quek *et al.*, 2018), and then focus HSF processing on the most relevant image locations such as the eye and mouth regions (Sergent, 1986; Sadr and Sinha, 2004; Blais *et al.*, 2008; Miellet, Caldara and Schyns, 2011; Rossion, 2014). LSF contrast is a reliable indicator towards the location of those regions, so favoring HSF processing within regions of high LSF contrast would allow for effective and resource-efficient processing of face identity. Such a process could in principle occur at any level of the visual hierarchy where receptive fields from various spatial frequency ranges are converging. However, to restrict HSF processing to relevant image regions via the use of feedback, at least the coarse retinotopic organization of the stimulus needs to be preserved in the feedback signal. Previous studies that explicitly investigated the spatial distribution of feedback effects to primary visual cortex however found evidence to the contrary. Williams et al. (2008) found feedback to foveal retinotopic areas of V1 to contain information about objects presented in the periphery. De Wit et al. (2012) found that V1 fMRI BOLD responses to bi-stable stimuli involve the entire V1, not only the retinotopic regions corresponding to the perceptual change. Even more strikingly, Hsieh et al. (2010) found evidence that the patterns of early visual cortex response (V1/V2/V3) to two-tone images were more similar to the patterns evoked by the same images’ grayscale photograph version if the participant knew the original image than it was to the identical but unrecognized two-tone image. Combined, these findings contradict a role for purely spatially defined feedback mechanisms.

Studies on location specific object coding in visual cortex however routinely find markers of enhanced object representation when the object appears in a retinotopic location that matches environmental statistics, such as e.g. a plane in the upper visual field, or a carpet in the lower visual field (Kravitz, Vinson and Baker, 2008; Kaiser and Haselhuhn, 2017; Kaiser, Moeskops and Cichy, 2018). These differences are typically found within the first 200 ms of visual processing (Kaiser, Moeskops and Cichy, 2018). Kok, Jehee, & De Lange (2012) argue that prior expectation of a relevant stimulus feature sharpens that features’ neural representation under noisy conditions. In line with this logic, suppressive feedback to regions of early visual cortex that do not match the retinotopic location of the presented stimulus could reduce stimulus ambiguity and would be measured as an overall decrease in response along with an enhancement of the stimulus-specific representation. While MEG recordings offer superior temporal resolution in comparison with fMRI recordings and could therefore potentially be suited to dissociate possible earlier retinotopic effects from possible later more global effects, they lack comparable spatial specificity and MEG responses obtained from traditional set-ups cannot currently be linked to a precise retinotopic target within early visual cortex (but see Iivanainen, Zetter and Parkkonen, 2020). To dissociate between retinotopic and more global effects, high spatial resolution fMRI should be combined with a temporally sensitive stimulus presentation paradigm such as temporal masking (e.g. Goffaux *et al.*, 2010). Alternatively, temporally sensitive recordings such as MEG or EEG could be combined with spatially distinct stimuli presentation methods such as foveal vs. peripheral stimulation to dissociate between potentially retinotopic transient feedback effects and later global effects.

### Cross-frequency, cross-region coupling

Since feedback signalling has been shown to be carried by neural oscillations in the alpha/beta range (Michalareas *et al.*, 2016), we expected to observe increased responses in these bands in inferior temporal or frontal regions during the processing of broadband intact compared to scrambled images. We further expected such alpha/beta band signals to correlate with the reduction of local processing markers in the gamma range during the processing of SF-redundant information (intact images) on a trial-by-trial basis. Our results do not support this hypothesis. While we do find descriptive condition differences in cross frequency power coupling between the Fusiform and Calcarine ROIs, the statistical evidence is comparatively weak (see figure 9 and supplementary figure 4). We found no significant differences between conditions for phase-amplitude coupling between any ROIs. While this could reflect a true lack of condition specific higher-level feedback to early visual cortex, several alternative explanations for the absence of strong evidence need to be considered. Instead of scanning the brain volume for regions that portray strong cross frequency coupling, we restricted our analysis to the ROIs we a-priory defined based on a combination of anatomical and functional criteria. The functional criterium was the maximum selective response to flickering images of intact faces in independent data. The fast-changing and transient presentation of each image, masked forward and backward by preceding and succeeding images, respectively, likely favoured feedforward responsive neural populations (Keysers *et al.*, 2001; Fahrenfort, Scholte and Lamme, 2007). It is well possible that cortical sources that provide feedback to early visual cortex when presented with face images are not the same that are most strongly, or most selectively activated during the processing of flickering face images. In this case, our ROI selection could not have optimally included potential sources of feedback. However, the limited spatial resolution of MEG makes it unlikely that the reconstructed source parcels would be spatially distinct for feedforward and feedback signals. More importantly, our (intact) stimuli were easily recognizable and thus not specifically designed to require strong reliance on top down context to be resolved (Parker and Krug, 2003; Parkkonen *et al.*, 2008).

Apart from ROI and stimulus selection, there are also methodological challenges to consider in determining cross frequency coupling. Detecting cross frequency coupling and unambiguously attributing it to a causal relationship between modulating and modulated frequency, is not a straightforward process. The isolation of the frequency bands in question (which are either chosen arbitrarily, based on previous literature or characteristics of (independent) data from the same participants, or some combination thereof) relies on frequency filters. Here, we chose zero-phase FIR filters with subsequent Hilbert transforms to extract instantaneous phase from the lower frequencies as well as power from the gamma range. The type of filter (causal vs. non-causal, filter order etc.) can have profound effects on the filtered signal (Widmann, Schröger and Maess, 2015). Further, the bandwidth of the filtered signal is crucially important for the detection of any cross frequency coupling (Berman *et al.*, 2012; Aru *et al.*, 2015). While the bandwidth of the modulating frequency signal can and should be narrow, especially when estimating instantaneous phase, the bandwidth of the modulated signal must be sufficiently wide to capture any modulations. Due to the Heisenberg-Gabor limit that inversely relates the precision with which a signal can be localized in time to the precision with which it can be localized in frequency, a signal that is modulated by a slower frequency (i.e., one with a longer phase interval, such as alpha) can be more sharply localized in the frequency domain than a signal modulated by a higher frequency, with a shorter phase interval (such as beta) (Dvorak and Fenton, 2014). Accordingly, we chose our gamma filter bandwidth to be sufficiently wide (twice the modulation frequency). Another dilemma in the estimation of coupling between two signals extracted from stimulus locked responses is non-stationarity. Non-stationary processes, such as most neurophysiological responses, contain spectral correlations across frequencies (Lii and Rosenblatt, 2002). Those correlations are easily misinterpreted as cross frequency coupling (Aru *et al.*, 2015). We only considered cross frequency coupling between distal regions making our results less likely to suffer from signal autocorrelations. It is however possible, that a common driving input caused spurious coupling. Finally, the (frequentist) statistical analysis of cross frequency coupling hinges on the appropriateness of the surrogate data used to estimate the null distribution. We here chose for a conservative approach where we create surrogate data by time-shifting the gamma power time-series by random offsets, within the same trial (Cohen, 2014). This disrupts any existing temporal relationship between the lower frequency and higher frequency time-series without removing potential overall power differences between conditions and trials and therefore provides a suitable null distribution. Prior to comparing between conditions on the group level, each individual participant’s cross frequency coupling was thus standardized by that same participant’s permutation distribution for each condition. Altogether, ROI selection that was not optimized for finding evidence of feedback signalling, short trials and conservative statistical choices might have obscured potential cross-frequency coupling effects in our design. While our primary aim here was to establish the effects of redundant image information on EVC processing load, future studies specifically designed to address feedback to EVC will be needed to investigate the causal mechanics of such LSF facilitation.

In conclusion, we find evidence for the LSF guidance of HSF image information that is expressed in both spatial frequency dominance patterns in early visual cortex and early visual cortex gamma. While our data does not provide evidence that higher level feedback from the selected regions of interest drives the exploitation of image information redundancies in early visual cortex, our findings are in line with coarse-to-fine theories of visual processing. Considering redundant information across spatial frequencies as an asset rather than a nuisance in visual signals could help to understand visual function as a flexible, recurrent process.

## Acknowledgments

We thank Marie Louise Holm Møller and Sigbjørn Hokland for their help with data acquisition. This work was supported by ERC Starting Grant 640448 to SSD, the Excellence of Science grant HUMVISCAT-30991544 awarded to VG as well as FNRS ASP grant 32704080 and FNRS travel grant awarded to KP. VG is supported by the Belgian national fund for scientific research (FRS-FNRS).

