## Supplemental figures for "Information redundancy across spatial scales modulates early visual cortical processing"

### Supplementary materials:

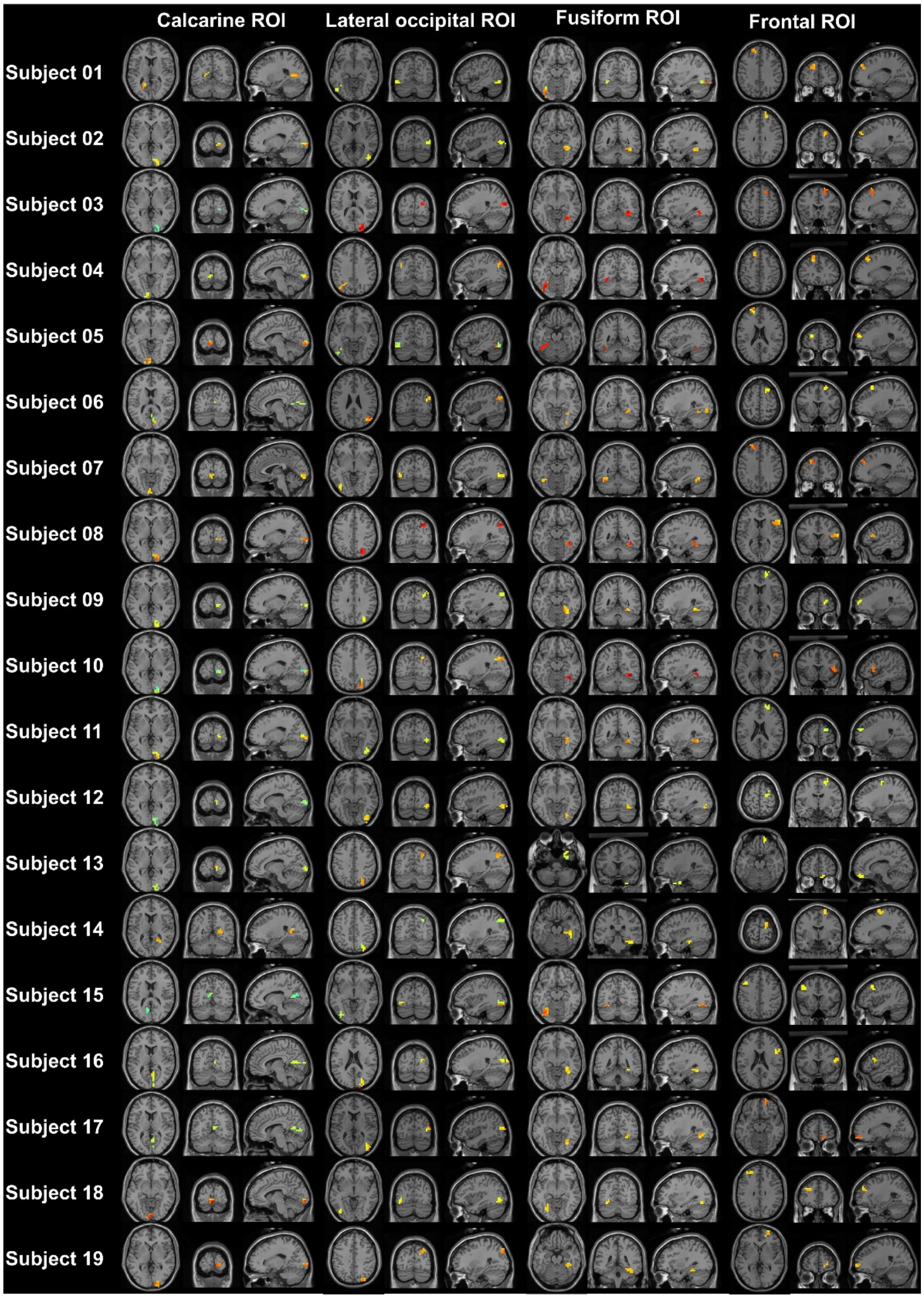


Figure 1: Source ROIs. Note that the lateralization of the face selective response in the Fusiform gyrus was used to select which hemisphere was used for further analysis.

Spatial frequency dominance classification (figure 5 in main manuscript) cannot be compared to a known ground truth. To demonstrate that source level classification can deliver robust results and generalizes across spatial frequencies, we constructed a classification problem with a known ground truth to evaluate our classification approach. We classified between intact and scrambled broadband image trials, and found the classifier's performance to be significantly above chance for all investigated ROIs (see Table 1).

| | BB INTACT VS. SCRAMBLED | GENERALIZATION TO LSF | GENERALIZATION TO HSF | DIFFERENCE IN GENERALIZATION PERFORMANCE (LSF-HSF) |
|---|---|---|---|---|
| **FUSIFORM** | $p<.05$, CS = $9.32x10^4$ | $p<.001$, CS = $3.89x10^4$ | $p<.001$, CS = $1.12x10^5$ | (1) $p<.05$, CS = $-1.26x10^4$ (2) $p<.05$, CS = $-4.29x10^3$ |
| **LATERAL OCCIPITAL** | $p<.05$, CS =$5.85x10^4$ | $p<.001$, CS =$6.25x10^4$ | $p<.001$, CS =$8.34x10^4$ | $p<.05$, CS =$-1.65x10^4$ |
| **FRONTAL** | (1) $p<.05$, CS = $6.25x10^3$ (2) $p<.05$, CS = $4.8x10^3$ | (1) $p<.05$, CS = $2.9x10^3$ (2) $p<.05$, CS = $2.82x10^3$ | $p<.001$, CS = 2.92x104 | $p<.05$, CS = $-1.24x10^4$ |
| **EVC** | $p<.001$, CS = 1.32x105 | $p<.001$, CS = $1.06x10^5$ | $p<.001$, CS = $1.63x10^5$ | $p<.001$, CS = $-3.56x10^4$ |

*Table 1: intact vs scrambled image classification results. LDA classifiers were trained on broadband trials to differentiate between intact and scrambled image trials. Classifiers generalized to both LSF and HSF conditions. CS = cluster statistic (maximum sum).*

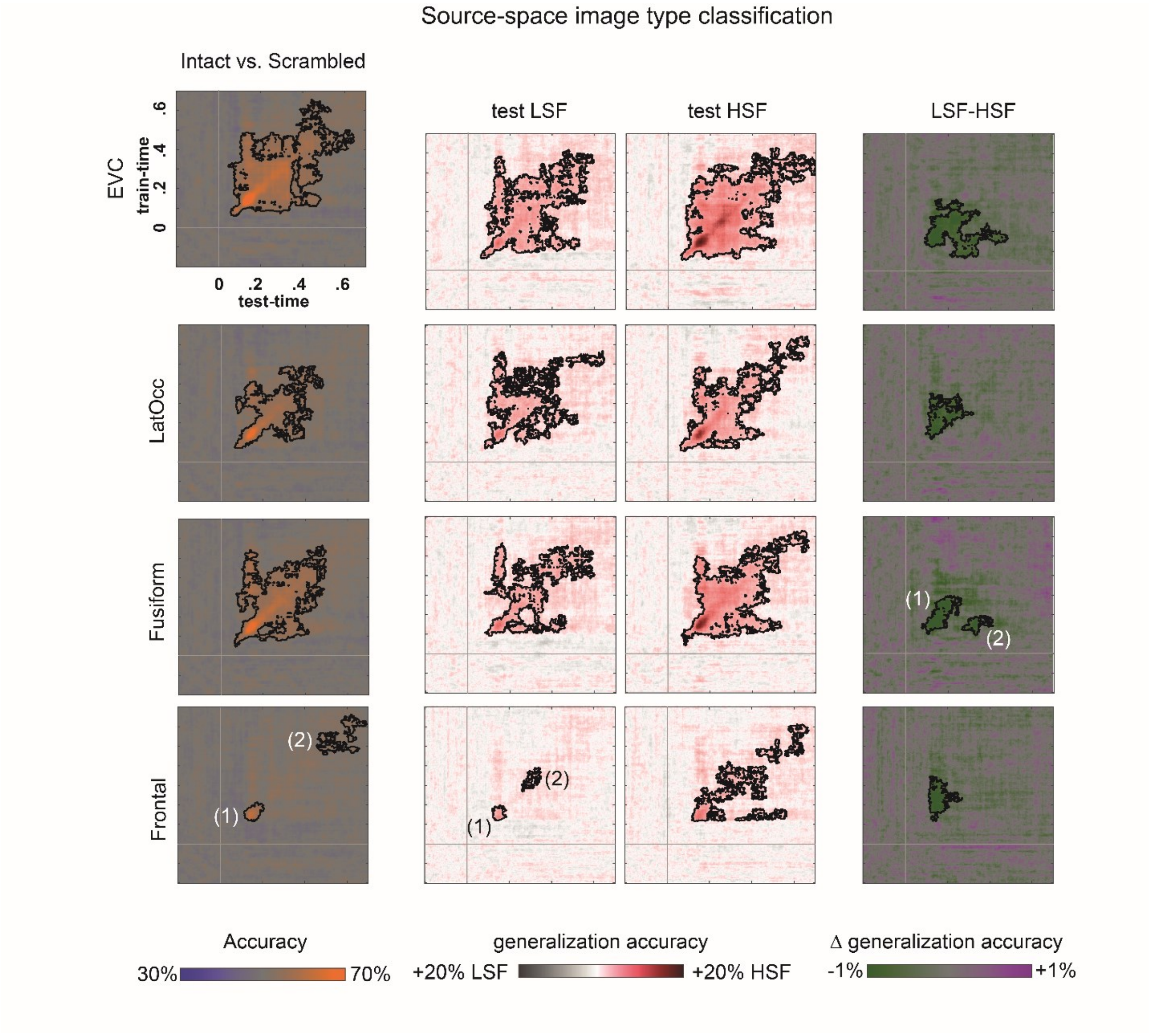


Figure 2: Classifiers trained on broadband image trial data generalized to both LSF and HSF trial data. Interestingly, generalization performance was significantly higher and widespread for HSF, compared to LSF image trials, indicating a preferential use of HSF information for image type classification.

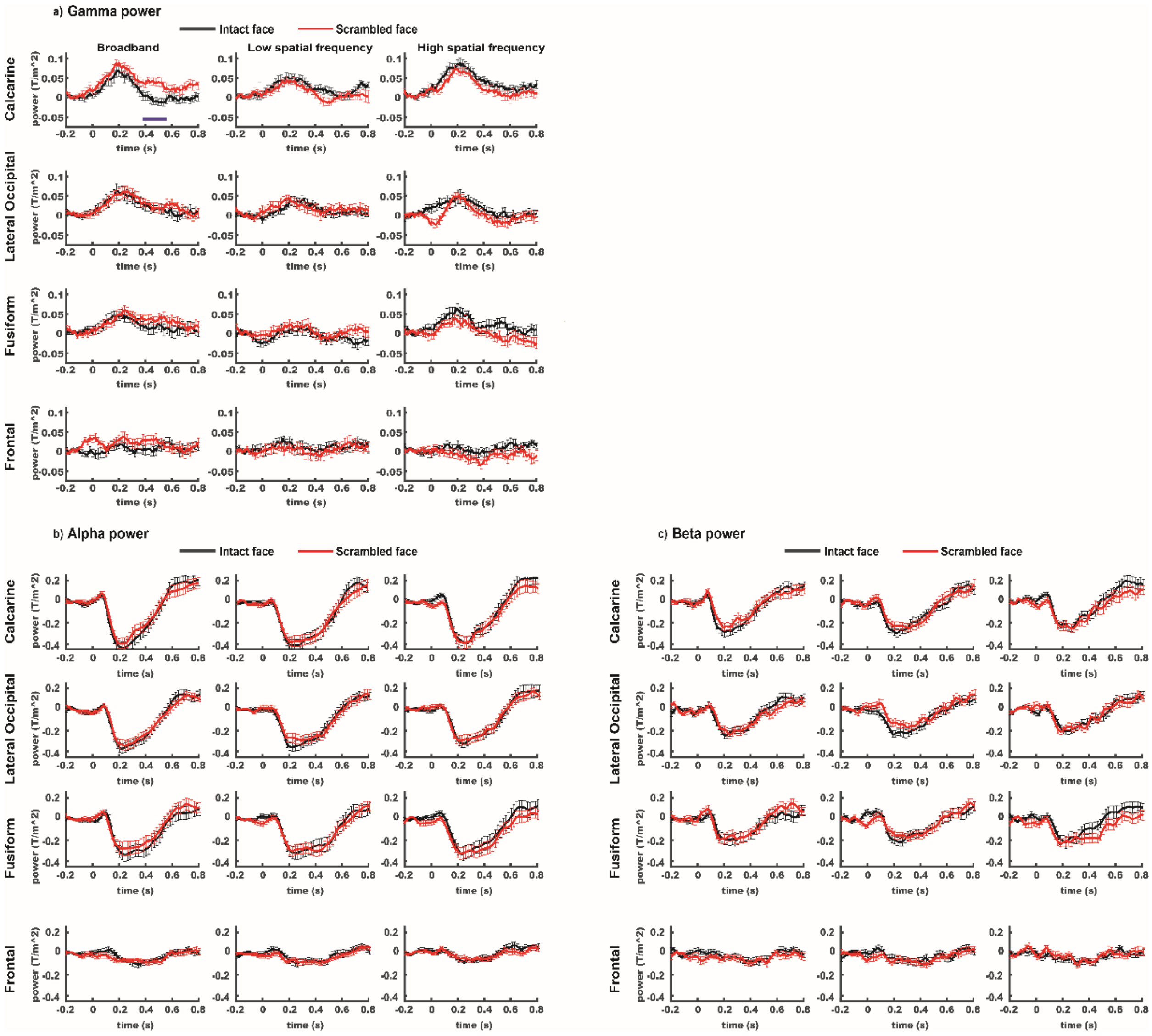


*Figure 3: Source power in the gamma (a), alpha (b) and beta (c) bands.  Error bars indicate standard error of the mean.*

#### *Lagged PPC between "higher level" ROI α/β and early visual cortex γ-power*

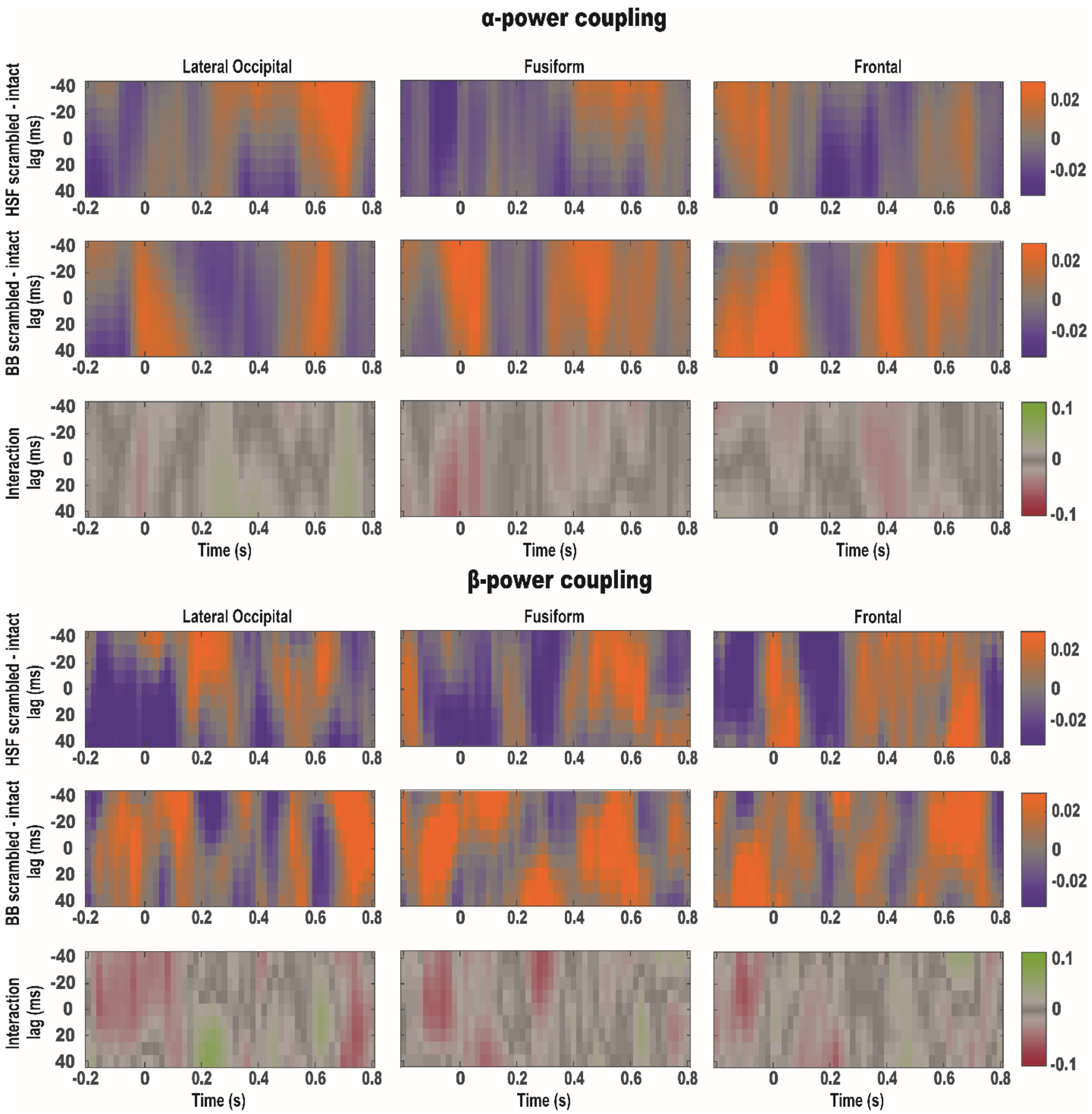


*Figure 4: Lagged PPC between "higher level" ROI α/β and early visual cortex γ-power. We tested for condition differences in coupling between high level ROI low frequency (α/β) band power and early visual cortex γ band power. To account for finite transmission velocities of cortical signals, we considered a ±40 ms lag. After correcting for multiple comparisons using cluster based permutation testing, none of the conditions differences or their interaction met our threshold for statistical significance (p<0.05).*
